# MitoHiFi: a python pipeline for mitochondrial genome assembly from PacBio High Fidelity reads

**DOI:** 10.1101/2022.12.23.521667

**Authors:** Marcela Uliano-Silva, João Gabriel R. N. Ferreira, Ksenia Krasheninnikova, Darwin Tree of Life Consortium, Giulio Formenti, Linelle Abueg, James Torrance, Eugene W. Myers, Richard Durbin, Mark Blaxter, Shane A. McCarthy

**Author notes:** correspondence, Marcela Uliano-Silva. contributed equally to this work (Marcela Uliano-Silva, João Gabriel R. N. Ferreira). João Gabriel R. N. Ferreira.

## Abstract

**Background:** PacBio high fidelity (HiFi) sequencing reads are both long (15-20 kb) and highly accurate (>Q20). Because of these properties, they have revolutionised genome assembly leading to more accurate and contiguous genomes. In eukaryotes the mitochondrial genome is sequenced alongside the nuclear genome often at very high coverage. A dedicated tool for mitochondrial genome assembly using HiFi reads is still missing.

**Results:** MitoHiFi was developed within the Darwin Tree of Life Project to assemble mitochondrial genomes from the HiFi reads generated for target species. The input for MitoHiFi is either the raw reads or the assembled contigs, and the tool outputs a mitochondrial genome sequence fasta file along with annotation of protein and RNA genes. Variants arising from heteroplasmy are assembled independently, and nuclear insertions of mitochondrial sequences are identified and not used in organellar genome assembly. MitoHiFi has been used to assemble 374 mitochondrial genomes (369 from 12 phyla and 39 orders of Metazoa and from 6 species of Fungi) for the Darwin Tree of Life Project, the Vertebrate Genomes Project and the Aquatic Symbiosis Genome Project. Inspection of 60 mitochondrial genomes assembled with MitoHiFi for species that already have reference sequences in public databases showed the widespread presence of previously unreported repeats.

**Conclusions:** MitoHiFi is able to assemble mitochondrial genomes from a wide phylogenetic range of taxa from Pacbio HiFi data. MitoHiFi is written in python and is freely available on github (https://github.com/marcelauliano/MitoHiFi). MitoHiFi is available with its dependencies as a singularity image on github (ghcr.io/marcelauliano/mitohifi:master).

## Introduction

Recent advances in genomics have opened the prospect of full genome sequencing of a wide range of species. Both global and taxonomically- or geographically-bounded projects have been initiated to exploit these new technologies to build reference genome libraries for all species on Earth. It is expected that these genome sequences will be a foundational dataset for new understanding in biology, new avenues in conservation and biodiversity monitoring, and new data to promote sustainable bioindustry [1,2]. The Earth Biogenome Project (EBP) [3] was founded to coordinate and promote biodiversity genomics through affiliated projects such as the Darwin Tree of Life project (DToL) [4], the Vertebrate Genomes Project (VGP) [5] and the Aquatic Symbiosis Genomics project (ASG) [6]. DToL is geographically-focussed, and aims to sequence all eukaryotic species living in and around the islands of Britain and Ireland. The goals of VGP are taxonomically-bounded (to sequence all vertebrate species), while those of ASG are defined by a particular biology (eukaryotes that live in symbiosis with prokaryotic or eukaryotic microbial partners). For each of these and other EBP projects, extensive development is needed in sample collection and preservation, nucleic acid extraction, sequencing, assembly, curation and annotation. A key driver in all these fields is the development of processes that can work at scale, processing hundreds to thousands of species rapidly.

One key advance has been the release of commercial long read sequencing platforms. Here we focus in particular on the PacBio zero mode waveguide, single molecule real time sequencing technology, and the particular deployment of this approach to produce long, high-quality reads using a circular consensus approach (called HiFi for high fidelity). These data have N50 read lengths of 15-20 kb, and accuracies Q20 or higher. This radically new data type has changed the landscape of genome assembly [7], and new toolkits (such as HiCanu [8] and HiFiAsm [9]) have been developed to exploit the very favourable characteristics of HiFi data.

Except for a few groups, all eukaryotes carry an essential cytoplasmic organelle, the mitochondrion. The mitochondrion derives from an ancient symbiosis between the last common ancestor of all eukaryotes and an alphaproteobacterial cell, and while many genes that were on the ancestral bacterial genome have been transferred to the nuclear genome, the mitochondrion generally retains a reduced, circular genome [10,11]. Mitochondria play a fundamental role in aerobic respiration and other key processes and the genes on the mitochondrial genome are essential for these functions [12]. While mitochondrial genome content varies extensively [13], in Metazoa the gene content of the mitochondrial replicon is fixed (11 or 12 protein coding genes, two ribosomal RNAs and a full set of tRNAs that decode the distinct organellar genetic code). Because the mitochondrial genome is haploid, and generally passes only through the maternal lineage, drift and linked selection tend to purge variants rapidly. Heteroplasmy, the presence of variant mitochondrial genomes within a single individual, has been thought to be generally rare and often associated with disease or normal somatic ageing processes. Repetitive sequence is thought to be generally absent from mitochondrial genomes. However analyses of long-read sequencing data identified frequent heteroplasmy and extended repeat content in vertebrate mitochondrial genomes [14]. These features were not evident in, or accessible through, previous short-read or PCR-and-sequence approaches.

The increase in the rate of generation of reference genomes across biodiversity using accurate long reads offers an opportunity to revisit the evolution of the gene content and structure of mitochondrial genomes. This opportunity requires development of robust pipelines that reliably assemble organellar genomes, correctly report repeat content, resolve heteroplasmy and distinguish true mitochondrial genome sequence from the frequent presence of nuclear genome inserted copies (nuclear mitochondrial transfers or NUMTs). Here we present MitoHiFi, software designed to use Pacbio HiFi reads to assemble and annotate complete mitochondrial genomes. MitoHiFi was developed within the Darwin Tree of Life Project and has been deployed to assemble mitochondrial genomes from hundreds of Metazoa and Fungi. MitoHiFi is distributed under the MIT licence, and is available on github (https://github.com/marcelauliano) and as a singularity image (ghcr.io/marcelauliano/mitohifi:master).

## Results

### MitoHiFi is a toolkit for robust assembly of mitochondrial genomes from HiFi data

MitoHiFi is written in python and orchestrates a series of external tools to select mitochondrial reads from whole genome sequencing datasets, filter out reads that are likely to derive from nuclear-mitochondrial transfers (NUMTs), assemble the genome, circularising it when possible and annotate it for protein and RNA genes. MitoHiFi also identifies possible heteroplasmic variants present in the data (Figure 1). We describe the processes embodied in MitoHiFi below.

**Figure 1:**
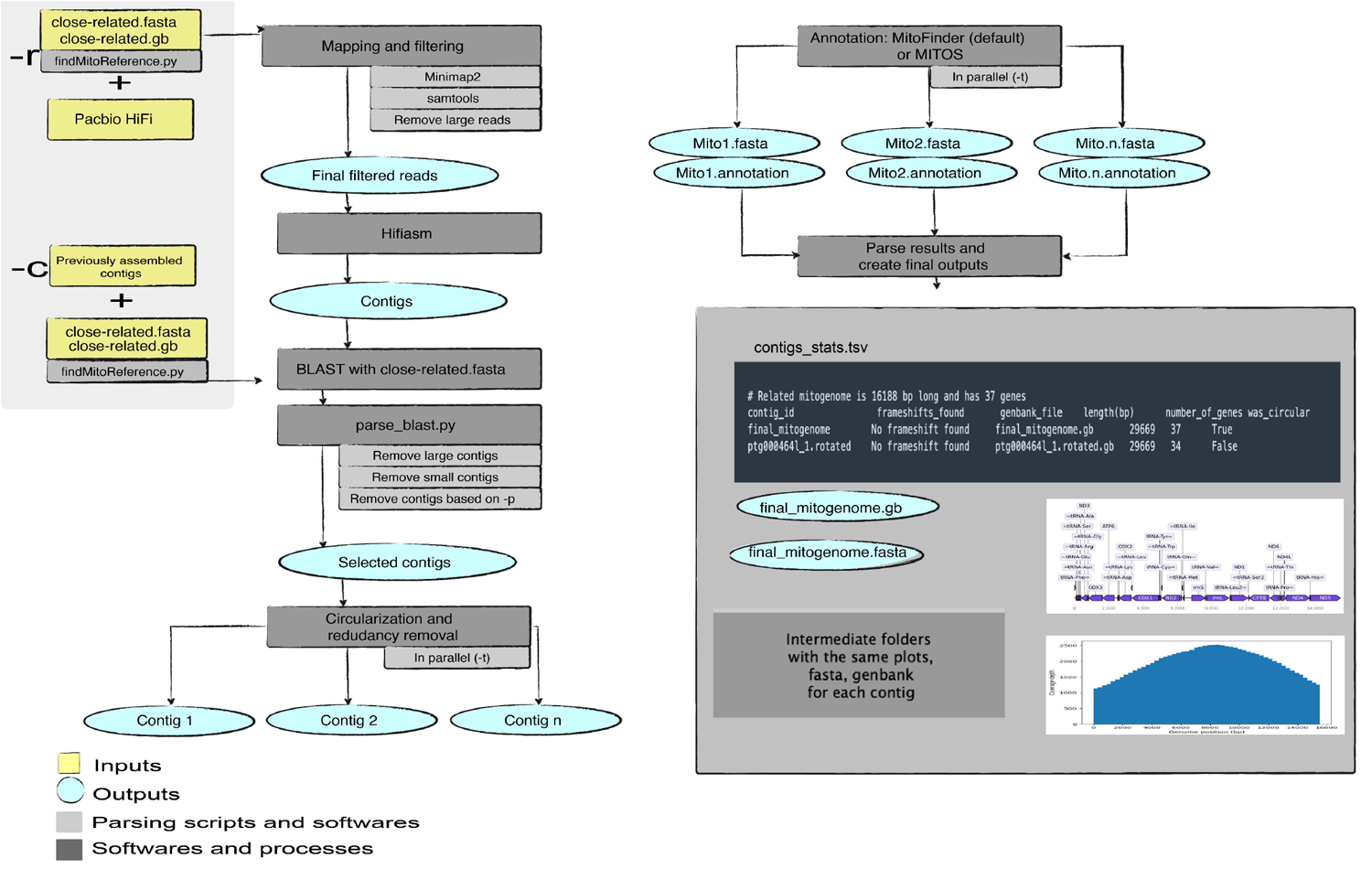
Outline of the MitoHiFi workflow. **A** Flow diagram of MitoHiFi processes. The inputs to MitoHiFi are a closely-related reference mitochondrial genome with either (i) raw Pacbio HiFi reads (`-r` parameter), or (ii) previously assembled contigs (`-c` parameter). **B** Outputs generated by MitoHiFi. A summary of the outputs generated by the pipeline, including the `contigs_stats.tsv` table summarising the metrics for the assembly, output sequence and annotation files, and graphical representations of coverage and annotation.

#### 1. Collecting PacBio HiFi reads based on reference mitochondrial genome(s) from closely-related species

As primary input, MitoHiFi takes Pacbio HiFi reads (`-r` flag), or contigs assembled from those reads (`-c`), and a “reference” mitochondrial genome from a closely-related species along with its annotation in GenBank format. The pipeline includes a script to automatically download closely-related mitochondrial genomes based on the NCBI taxonomy (`findMitoReference.py`). The defaults are set to download a single reference genome from the same or most-closely related species, but multiple references can be downloaded by setting `-n` (>1). When MitoHiFi is starting from reads (`-r`), it first maps the reads to this reference mitochondrial genome. The aligned readset is filtered to exclude reads that have lengths more than the exact length of the reference genome, as these are likely to derive from NUMT insertions [9]. However, this parameter can be changed when appropriate (flag `--max-read-len`).

#### 2. Assembly of mitochondrial reads and filtering of contigs to avoid NUMTs

The filtered readset is then assembled with Hifiasm [9] and assembled contigs are parsed through a series of filters to yield final mitochondrial genome candidates. The contigs are compared to the reference mitochondrial genome using BLAST+ [15]. Contigs are retained if

i. over 50% of their sequence length is present in the BLAST match with the reference (the user can change this threshold with the flag *-p*).
ii. the contig is less than five times the length of the reference (long contigs are likely to be NUMTs).
iii. the contig is >80% of the length of the reference (smaller contigs are likely to be incomplete).

All BLAST parsing information is saved in intermediate files *parsed_blast*.*txt* and *parsed_blast_all*.*txt* (see github page for intermediate output column details).

#### 3. Genome circularization

All candidate contigs are processed through circularizationCheck [16]. This module uses self-BLAST of each contig to identify sequence redundancy between its ends. As default >220 bp of overlap is required but the user can change this by setting `--circular-size` to a required length. MitoHiFi outputs circularisation information to *all_contigs*.*circularisationCheck*.*txt*.

#### 4. Annotating mitochondrial genomes

Candidate contigs are annotated in parallel (`-t` flag for number of threads) using MitoFinder (the default) [16] or MITOS [17] (using flag `--mitos`). The mitochondrial genetic code should be set with the flag `-o`. MitoFinder finds protein coding genes through BLAST similarity searches using the reference nucleotide and protein sequences. MITOS performs *de novo* annotation based on its database of orthologous genes. Both annotation pipelines use mitfi [18] to identify and classify tRNAs. MitoHiFi outputs the annotation for all assembled contigs in multiple formats (GenBank [gb], general feature format [gff], FASTA) and produces plots of the annotated features and of reads mapping coverage, if initiated from reads (`-r`). After annotation, MitoHiFi identifies the tRNA-Phe locus and rotates the mitochondrial genome to have its start at the first base of this locus, following established convention. If no contig has a tRNA-Phe locus, then the most frequent tRNA locus among all contigs is chosen as reference for rotation.

#### 5. Choosing a representative mitochondrial genome

We find that MitoHiFi frequently assembles more than one candidate mitochondrial genome from raw read data. This may be due to read error and high coverage, or true heteroplasmy (see Discussion). MitoHiFi outputs a sequence file (FASTA format) and annotation (GenBank format) for each contig, and selects a final representative mitochondrial genome using the following criteria:

i. all potential contigs are sorted by number of genes annotated in relation to the reference, then:
  a. it searches for contigs that were classified as circular (and has had sequence redundancy removed)
  b. have a similar size to the reference mitochondrial size
  c. and its annotation includes no genes that contain a frameshift.
ii. if no contig passes the criteria in I, MitoHiFi will select the circular contig that follows at least two of the criteria above in order A>B>C.

The selected main mitochondrial genome assembly files are renamed *final_mitogenome*.*fasta* and *final_mitogenome*.*gb*, and a graphical representation of the annotation is produced (*final_mitogenome*.*annotation*.*png*) along with a plot of the reads (`gbk.HiFiMapped.bam.filtered.fasta`) mapped to it (*final_mitogenome*.*coverage*.*png*). Mapping quality filtering for the final coverage plots can be set with the flag `-covMap`. MitoHiFi also outputs annotation and coverage plots for all other potential contigs (*contigs_annotations*.*png*, *coverage_plot*.*png*). To produce all the potential contigs coverage plot, they are concatenated in a file to use all as reference for mapping. Coverage plots are only produced when MitoHiFi is started with the flag `-r`.

MitoHiFi also produces a summary file *contigs_stats*.*tsv* that reports which closely-related reference was used and gives details of each assembled mitochondrial genome (including assembly size, number of genes annotated, presence of frameshifts in genes and circularisation data) (Figure 1B). Finally, MitoHiFi produces a *shared_genes*.*tsv* file that presents a comparison of genes annotated in the reference in relation to genes annotated in each potential contigs (including final) for a quick inspection of the annotation.

#### 6. Using MitoHiFi for mitochondrial genomes from non-metazoan species and for plastid genomes

Metazoan mitochondrial genomes are generally relatively small and have limited gene content. An exception is found in Cnidaria, where gene content and size can be larger. MitoHiFi performs well with these genomes as well. For other lineages, where gene content and size can vary greatly even between closely related taxa, MitoHiFi offers alternate approaches. For the assembly of mitochondrial genomes from Viridiplantae (plants) the parameter `-a plant` should be called. This will prevent `parse_blast.py` from filtering out large contigs as plant mitochondrial genomes can greatly vary in size even between closely-related taxa [19]. Plastid genomes are similarly variable in size and content. For plastid genome assembly `findMitoReference.py` should be run with the flag `–type chloroplast` to find a closely-related plastid genome. While MitoFinder is used as default for annotation, if the user wishes to use MITOS for fungal genomes, the parameter `-a fungi` should be used.

### Assembly of mitochondrial genomes at scale

MitoHiFi has been deployed to assemble mitochondrial genomes from species analysed by DToL (350 species to December 2022), the VGP (22 species), and ASG (2 species) from 12 phyla and 39 orders of Metazoa and Fungi (Figure 2). The assemblies have been submitted to the International Nucleotide Sequence (INSDC) databases (ENA, NCBI, DDBJ) or to the VGP GenomeArk and a list of accession numbers can be found in Supplementary Table 1.

**Figure 2:**
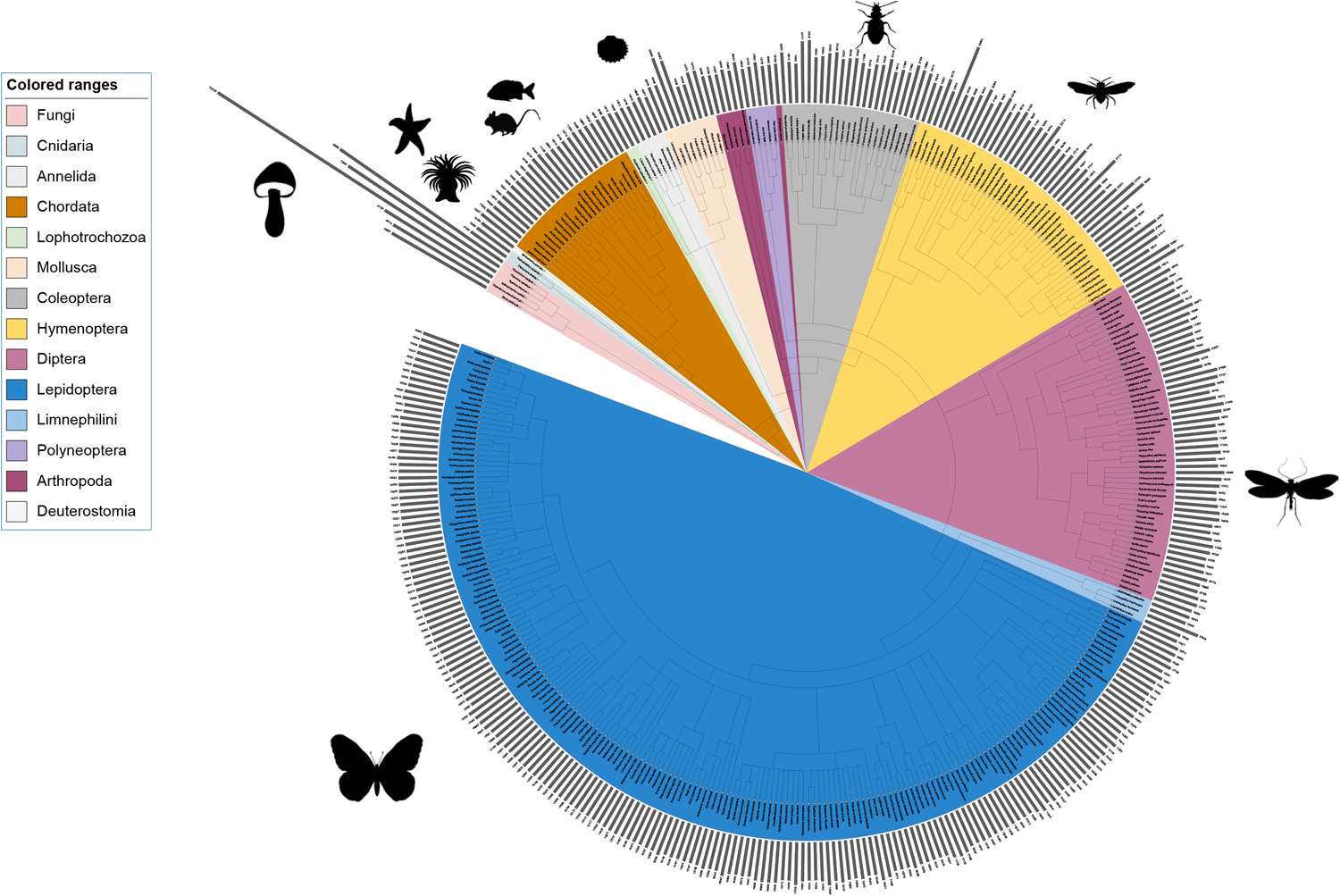
Mitochondrial genomes assembled with MitoHiFi for DToL, ASG and VGP species. The 374 mitochondrial genomes assembled using MitoHiFi are shown. The tree is derived from NCBI TaxonomyDB [20,21], and was visualised in iToL [22]. The histogram shows the mitochondrial genome length for each species. Some species icons are from [PhyloPics2.0](https://beta.phylopic.org/)

### MitoHiFi identifies additional repeat content in previously published genomes

Sixty of the species assembled here with MitoHiFi had mitochondrial genomes previously sequenced and submitted to the INSDC databases (Supplementary Table 2, Figure 3A). To assess the performance of MitoHiFi we compared the new MitoHiFi assemblies to these published assemblies (Figure 3). The majority of species had assemblies of a similar size and with a nucleotide identity above 96%. Only three MitoHiFi assemblies were smaller than the previously assembled reference. The *Flammulina velutipes* (Fungi, Basidiomycota, Agaricomycetes) MitoHiFi assembly was 8,927 bp smaller than the database reference JN190940.1. Apart from that 8kb portion, both assemblies have the same t-RNAS, rRNAs and protein coding genes annotated and a nucleotide similarity of 97%. We investigated the 8kb sequence unique to JN190940.1 through mapping our Pacbio HiFi reads to it, and found no evidence of reads spanning that sequence. We also compared this additional sequence using BLAST [15] against the NCBI nucleotide database and found no evidence of it being present in other fungal mitochondrial genomes. The additional segment in JN190940.1 contains no essential conserved genes. The difference between the MitoHiFi assembly and the published one could be technical due to accidental inclusion of a segment due to the technology used in cloning and PCR or true biological variation between isolates.

**Figure 3:**
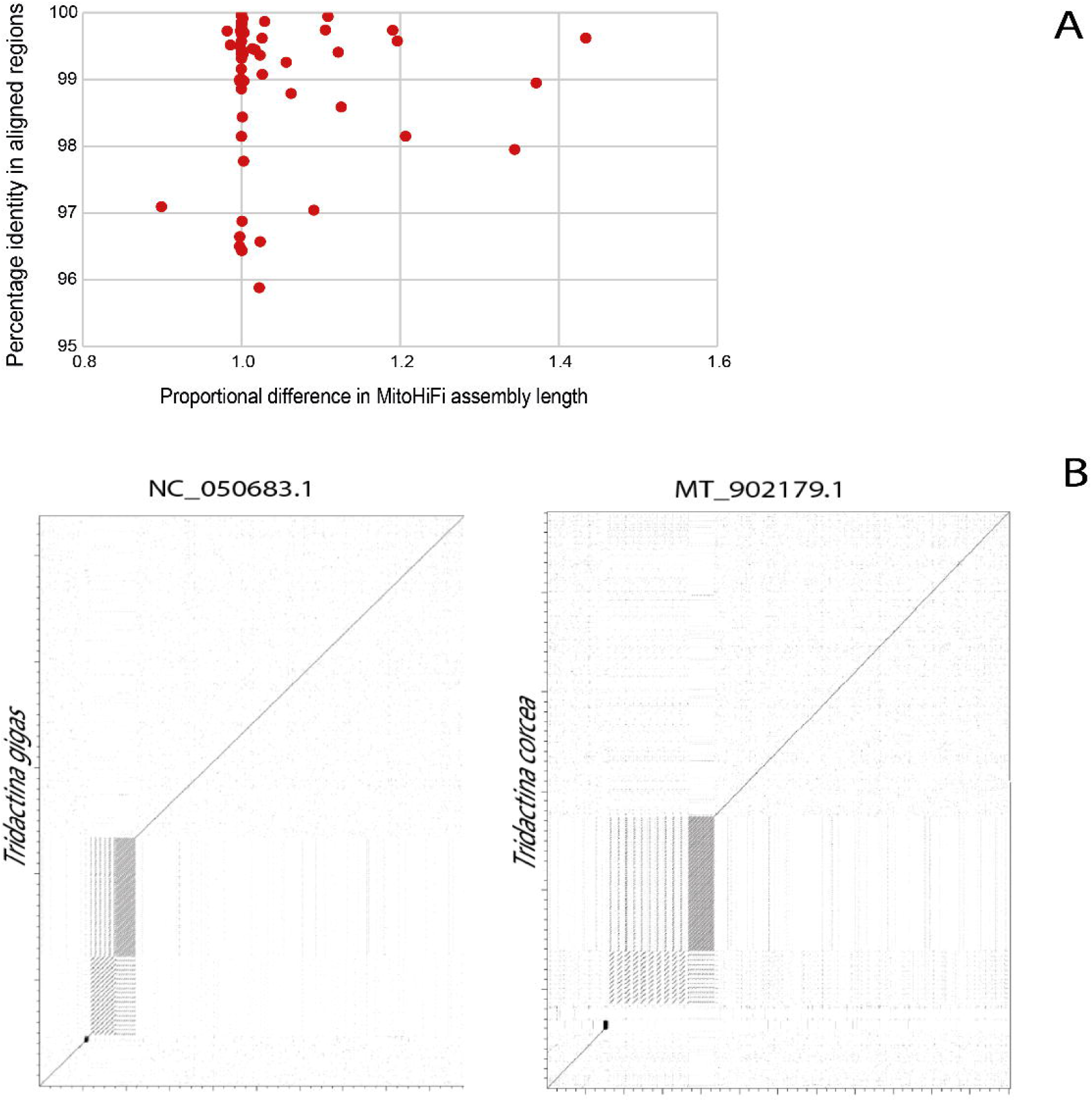
MitoHiFi assemblies often include sequence absent from previous mitochondrial genome assemblies. **A** For 60 species, the new MitoHiFi assemblies were compared to sequences available for the same species in INSDC databases. For each pair the percent nucleotide identity of the aligned sequences and the proportional size of the MitoHiFi assembly were measured with FastANI. **B** Comparison (nucmer) of the sequences of the MitoHiFi (y-axes) and previously published mitochondrial genomes (x-axes) of the giant clams *Tridacna gigas* and *Tridacna crocea*. The MitoHiFi assemblies include multiple copies of repeat sequences, supported by individual long reads, that are likely collapsed in the previously published sequences.

Sixteen mitochondrial genomes assembled by MitoHiFi were substantially larger than their previous version. Alignment and gene content investigations show that repeats are the cause of longer MitoHiFi assemblies (Figure 3B, Supplementary Figure 1). A dotplot and a nucleotide inspection of *Tridacna gigas* (Metazoa, Mollusca, Bivalvia) assemblies showed three types of repeats that are present in both assemblies, but those repeats have more copies present in the assembly built by MitoHiFi and are supported by HiFi data. The same is true for *Tridacna corsea* (Figure 3B). Other dotplots are shown in Supplementary Figure 1. For these 16 assemblies, where MitoHiFi genomes are larger than the previous reference, the percentage of nucleotide identity in the aligned portions is always above 95%, making it unlikely that MitoHiFi is including NUMTS in the final assembled sequence.

### Assembling non-metazoan mitochondrial and plastid genomes

Although MitoHiFi was developed and optimised for the assembly of metazoan mitochondrial genomes, which have a limited gene content and relatively small span, it successfully assembled mitochondria from other taxa, and other organellar genomes. For example, we successfully assembled fungal mitochondrial genomes from 43 to 133 kb (Figure 2). Plant organellar genomes present two challenges: high levels of heteroplasmy and presence of long repeat sections. Currently we recommend that plant organellar genome estimates generated by MitoHiFi should always be checked and manually finished.

## Discussion

In the emerging era of reference genomics, where chromosomally-complete assemblies are sought, PacBio HiFi reads are a key data type [7]. Because of their length and quality, HiFi data support rapid and robust assembly of any genome, including those of organelles. Existing organelle genome assembly toolkits ([16,23]) were designed for short reads, and deal efficiently with their limitations. However short reads cannot resolve tandem and other repeats and may not be able to distinguish mitochondrial insertions in the nuclear genome from true organellar sequence. We have built MitoHiFi to best make use of long, accurate HiFi reads. MitoHiFi analysis can be initiated from raw HiFi data or from reads pre-assembled into primary contigs, for example using Hifiasm [9] or HiCanu [8]. However, these assemblers are not designed to assemble organellar genomes, which will be present at high relative coverage compared to the target nuclear genome, and usually emit assembly estimates that include a multiplicity of subgenomes and misassemblies or reject the mitochondria as being a multicopy repeat.

MitoHiFi performs best when using raw read data, as this allows the tool to effectively avoid reads derived from NUMTs (and equivalent plastid transfers, NUPTs) and to identify likely assembled NUMT/NUPT loci using length cutoffs. MitoHiFi uses a relatively generous length cutoff because historical use of short read or PCR-based methods to retrieve organellar genomes may have resulted in collapse of repeat regions and thus underestimation of true genome length. In our analyses we identified many examples where the previously published assemblies differed from MitoHiFi assemblies because of a relative lack of repeat sequences in the published assemblies (Figure 3B, Supplementary Figure 1). Inspection of MitoHiFi outputs shows that these filters are effective in avoiding NUMTs/NUPTs (see Supplementary Figure 2). We note that the toolkit also emits these “other” assemblies, and would advise that if the final assembly generated for a species does not fit within expected parameters (presenting a small number of genes, very large repeats present or exhibiting many gene frameshifts), further manual validation should be performed.

The choice of the final reference assembly is based on comparison to previously assembled genomes available in public databases. The default parameter for coverage is set to 50% (*i*.*e*., at least 50% of the sequence of the contig has to be present in the BLAST match with the closely-related species mitochondrial genome), but this may be raised as high as 90% where the species being analysed is part of a richly-sampled group with highly constrained genome content and structure, such as Vertebrata. However, for other taxa such as Hymenoptera and Mollusca, the 50% match length cutoff is required (and indeed may be too strict) because these clades have mitochondrial genomes with highly variable gene orders [24] and a diversity of repeat lengths and copy numbers. For taxa with known variability, it is recommended that the pipeline is run using multiple different references (using the `findMitoReference.py -n` flag) and exploring different match length proportions.

MitoHiFi incorporates two tools for protein coding gene finding and annotation, MitoFinder (the default) [16] and MITOS (flag *--mitos*) [17]. Both annotators use mitfi [18] to identify tRNAs as default. Automatic annotation remains a challenging task, particularly for organellar genomes that contain genes not in the core set, and where intron splicing of RNA and protein coding genes is common. Because of this, annotations produced by MitoHiFi should be checked manually before submission to databases.

Although MitoHiFi was optimised to assemble metazoan mitochondrial genomes, it has performed well with Fungi where the mitochondrial genomes assembled ranged from 43 to 133 kb (Figure 2). MitoHiFi can be used to identify plant mitochondria and chloroplast genomes, but we have observed that the final sequences for these genomes are often incomplete. While MitoHiFi parameters can be adjusted to reflect the large variability in size and gene content of plant mitochondrial and plastid genomes, the presence of large repeat segments in mitochondrial and plastid genomes and high heteroplasmy makes current assembly outputs less than optimal. The plastid genome is characterised by the presence of a large inverted repeat that carries the small and large subunit ribosomal RNA genes (and sometimes other loci) [25]. This repeat can be longer than a single HiFi read, and it is difficult to fully resolve the orientation of the two non repeat segments. In practice, the plastid genome is usually present as two isomers generated by inversion across the repeats, and thus linear representations of plastid assemblies must select one of the two possible paths through the repeats. Graph representation of these genomes would better represent the actual structures found. Plant mitochondrial genomes are also found as multiple isomers, and the sequence can contain several repeat segments of varying lengths. Again HiFi reads may not span these repeats, and the only way to represent the genome fully would be through a graph structure. We are working on methods to use graph representations of plastid genome assembly to emit resolved structures of the different organellar genome populations present in plants.

## Conclusions

MitoHiFi efficiently assembles and annotates mitochondrial genomes using PacBio HiFi reads. We have used it to assemble 374 mitochondrial genomes from major biodiversity genomics projects. MitoHiFi is openly available on github as pure python code and as a docker container under the MIT licence.

## Methods

### Species analysed

The data analysed here were generated as part of the DToL, VGP and ASG projects. Each of the target species was sequenced to ∼25 fold coverage in PacBio HiFi reads of the nuclear genome following standard protocols. The nuclear genomes have been assembled using HiFi and Illumina Hi-C data, and will be reported elsewhere. Supplementary Table 1 presents the INSDC ascension numbers or VGP links for the genome sequences of each species analysed.

### MitoHiFi pipeline

MitoHiFi was written in python3. It incorporates other tools as shown in Figure 1. MitoHiFi was run from reads (-r) with default parameters for all species presented here apart from some fungi species. For *Mucor piriformis, Flammulina velutipes, Pleurotus ostreatus* and *Agaricus bisporus*, reads were first assembled with MBG ([26] parameters: -k 1001 -w 250 -a 5 -u 150) to obtain contigs that were then input to MitoHiFi with the `-c` flag. The github page can be accessed for detailed documentation on all final and intermediate outputs by MitoHiFi.

### Comparative analyses

Sixty mitochondrial genomes assembled by MitoHiFi (Supplementary Table 2) were compared with their previous reference found on INSDC with FastANI [27] for nucleotide identity and dotter [28] (Figure 3 and Supplementary Figure 1).

### Graphical representations

The tree in Figure 2 was derived from the TaxonomyDB tree using ETE based on the NCBI TaxIDs for each species. The figure was generated in ITOL [22]. Further modification, including addition of some species silhouettes from [PhyloPics2.0](https://beta.phylopic.org/) and others was performed in Adobe Illustrator.

### Data and software Accessibility

Supplementary Table 1 presents the INSDC accession numbers of the genome sequences of all the species analysed. The mitochondrial genomes have been submitted along with the nuclear genome sequences. MitoHiFi is available on github (https://github.com/marcelauliano/MitoHiFi) and as a docker image ghcr.io/marcelauliano/mitohifi:master) under the MIT licence.

