## SupplementaryTable1 for "MitoHiFi: a python pipeline for mitochondrial genome assembly from PacBio High Fidelity reads"

| **Supplementary Table 1:** Species from Figure 2 and their as | **Species name** | **NCBI Accession number** |
|---|---|---|
| | Acipenser ruthenus | GCA 902713425.1 |
| | Accipiter gentilis | GCA 929443795.1 |
| | Marthasterias glacialis | GCA 911728455.1 |
| | Pholis gunnellus | GCA 910591455.1 |
| | Taurulus bubalis | GCA 910589615.1 |
| | Apoderus coryli | GCA 911728435.1 |
| | Cantharis rustica | GCA 911387805.1 |
| | Coccinella septempunctata | GCA 907165205.1 |
| | Harmonia axyridis | GCA 914767665.1 |
| | Malachius bipustulatus | GCA 910589415.1 |
| | Ocypus olens | GCA 910593695.1 |
| | Pterostichus madidus | GCA 911728475.1 |
| | Pyrochroa serraticornis | GCA 905333025.1 |
| | Rhagonycha fulva | GCA 905340355.1 |
| | Bibio marci | GCA 910594885.1 |
| | Bellardia pandia | GCA 916048285.1 |
| | Cheilosia vulpina | GCA 916610125.1 |
| | Chrysotoxum bicinctum | GCA 911387755.1 |
| | Coremacera marginata | GCA 914767935.1 |
| | Eristalis arbustorum | GCA 916610145.1 |
| | Eristalis pertinax | GCA 907269125.1 |
| | Eristalis tenax | GCA 905231855.1 |
| | Gymnosoma rotundatum | GCA 916610165.1 |
| | Melanostoma mellinum | GCA 914767635.1 |
| | Sicus ferrugineus | GCA 922984085.1 |
| | Tachina fera | GCA 905220375.1 |
| | Xanthogramma pedissequum | GCA 910595825.1 |
| | Acanthosoma haemorrhoidale | GCA 930367205.1 |
| | Aelia acuminata | GCA 911387785.1 |
| | Abrostola tripartita | GCA 905340225.1 |
| | Acronicta aceris | GCA 910591435.1 |
| | Inachis io | GCA 905147045.1 |
| | Nymphalis urticae | GCA 905147175.1 |
| | Amphipyra berbera | GCA 910594945.1 |
| | Aporia crataegi | GCA 912999735.1 |
| | Autographa gamma | GCA 905146925.1 |
| | Biston betularia | GCA 905404145.1 |
| | Boloria selene | GCA 905231865.2 |
| | Catocala fraxini | GCA 930367265.1 |
| | Celastrina argiolus | GCA 905187575.1 |

| | | |
|---|---|---|
| | Colias croceus | GCA 905220415.1 |
| | Cosmia trapezina | GCA 905163495.1 |
| | Deilephila porcellus | GCA 905220455.1 |
| | Euproctis similis | GCA 905147225.1 |
| | Glaucopsyche alexis | GCA 905404095.1 |
| | Gymnoscelis rufifasciata | GCA 929108375.1 |
| | Hydraecia micacea | GCA 914767645.1 |
| | Hypena proboscidalis | GCA 905147285.1 |
| | Laothoe populi | GCA 905220505.1 |
| | Luperina testacea | GCA 927399505.1 |
| | Lycaena phlaeas | GCA 905333005.1 |
| | Lymantria monacha | GCA 905163515.1 |
| | Maniola jurtina | GCA 905333055.1 |
| | Mellicta athalia | GCA 905220545.1 |
| | Melitaea cinxia | GCA 905220565.1 |
| | Melanargia galathea | GCA 920104075.1 |
| | Mimas tiliae | GCA 905332985.1 |
| | Mythimna ferrago | GCA 910589285.1 |
| | Noctua fimbriata | GCA 905163415.1 |
| | Noctua pronuba | GCA 905220335.1 |
| | Notodonta dromedarius | GCA 905147325.1 |
| | Notocelia uddmanniana | GCA 905163555.1 |
| | Notodonta ziczac | GCA 918843915.1 |
| | Nymphalis polychloros | GCA 905220585.1 |
| | Pararge aegeria | GCA 905163445.1 |
| | Parapoynx stratiotata | GCA 910589355.1 |
| | Phalera bucephala | GCA 905147815.2 |
| | Pheosia tremula | GCA 905333125.1 |
| | Phlogophora meticulosa | GCA 905147745.1 |
| | Pieris brassicae | GCA 905147105.1 |
| | Pieris napi | GCA 905475465.1 |
| | Pieris rapae | GCA 905147795.1 |
| | Pyrgus malvae | GCA 911387765.1 |
| | Spilosoma lubricipeda | GCA 905220595.1 |
| | Thyatira batis | GCA 905147785.1 |
| | Thymelicus sylvestris | GCA 911387775.1 |
| | Tinea trinotella | GCA 905220615.1 |
| | Vanessa atalanta | GCA 905147765.2 |
| | Vanessa cardui | GCA 905220365.1 |
| | Xestia xanthographa | GCA 905147715.1 |
| | Chrysoperla carnea | GCA 905475395.1 |

| | | |
|---|---|---|
| | Ischnura elegans | GCA 921293095.1 |
| | Nemurella pictetii | GCA 921293315.1 |
| | Anoplius nigerrimus | GCA 914767735.1 |
| | Bombus hortorum | GCA 905332935.1 |
| | Cerceris rybyensis | GCA 910591515.1 |
| | Dolichovespula sylvestris | GCA 918808275.1 |
| | Ichneumon xanthorius | GCA 917499995.1 |
| | Lasioglossum lativentre | GCA 916610255.1 |
| | Macropis europaea | GCA 916610135.1 |
| | Seladonia tumulorum | GCA 913789895.1 |
| | Sphecodes monilicornis | GCA 913789915.1 |
| | Tenthredo notha | GCA 914767705.1 |
| | Vespa crabro | GCA 910589235.1 |
| | Vespula germanica | GCA 905340365.1 |
| | Vespula vulgaris | GCA 905475345.1 |
| | Diadumene lineata | GCA 918843875.1 |
| | Aplidium turbinatum | GCA 918807975.1 |
| | Canis lupus | GCA 905319855.2 |
| | Cervus elaphus | GCA 910594005.1 |
| | Rattus norvegicus | GCA 015227675.2 |
| | Lineus longissimus | GCA 910592395.1 |
| | Gari tellinella | GCA 922989275.1 |
| | Patella pellucida | GCA 917208275.1 |
| | Steromphala cineraria | GCA 916613615.1 |
| | Clusia tigrina | GCA 920105625.1 |
| | Criorhina berberina | GCA 917880715.1 |
| | Eupeodes latifasciatus | GCA 920104205.1 |
| | Platycheirus albimanus | GCA 916050605.1 |
| | Scaeva pyrastri | GCA 905146935.1 |
| | Syritta pipiens | GCA 905187475.1 |
| | Thecocarcelia acutangulata | GCA 914767995.1 |
| | Volucella inanis | GCA 907269105.1 |
| | Xylota sylvarum | GCA 905220385.1 |
| | Limnephilus lunatus | GCA 917563855.1 |
| | Limnephilus marmoratus | GCA 917880885.1 |
| | Acleris sparsana | GCA 923062465.1 |
| | Agonopterix subpropinquella | GCA 922987775.1 |
| | Agriopis aurantiaria | GCA 914767915.1 |
| | Agrochola circellaris | GCA 914767755.1 |
| | Agrochola macilenta | GCA 916701695.1 |
| | Amphipyra tragopoginis | GCA 905220435.1 |

| | | |
|---|---|---|
| | Anthocharis cardamines | GCA 905404175.1 |
| | Apamea monoglypha | GCA 911387795.1 |
| | Aplocera efformata | GCA 921293045.1 |
| | Apotomis turbidana | GCA 905147355.1 |
| | Aricia agestis | GCA 905147365.1 |
| | Atethmia centrago | GCA 905333075.2 |
| | Autographa pulchrina | GCA 905475315.1 |
| | Bembecia ichneumoniformis | GCA 910589475.1 |
| | Blastobasis adustella | GCA 907269095.1 |
| | Blastobasis lacticolella | GCA 905147135.1 |
| | Campaea margaritaria | GCA 912999815.1 |
| | Carcina quercana | GCA 910589575.1 |
| | Chrysoteuchia culmella | GCA 910589605.1 |
| | Clostera curtula | GCA 905475355.1 |
| | Craniophora ligustri | GCA 905163465.1 |
| | Crocallis elinguaria | GCA 907269065.1 |
| | Cyaniris semiargus | GCA 905187585.1 |
| | Cydia splendana | GCA 910591565.1 |
| | Dryobotodes eremita | GCA 917490735.1 |
| | Eilema depressum | GCA 914767945.1 |
| | Eilema sororcula | GCA 914829495.1 |
| | Emmelina monodactyla | GCA 916618145.1 |
| | Endotricha flammealis | GCA 905163395.1 |
| | Ennomos fuscantarius | GCA 905220475.1 |
| | Ennomos quercinarius | GCA 910589525.1 |
| | Erannis defoliaria | GCA 905404285.1 |
| | Erebia aethiops | GCA 923060345.1 |
| | Erebia ligea | GCA 917051295.1 |
| | Erynnis tages | GCA 905147235.1 |
| | Eulithis prunata | GCA 918843925.1 |
| | Euplexia lucipara | GCA 921972225.1 |
| | Eupsilia transversa | GCA 914767815.1 |
| | Fabriciana adippe | GCA 905404265.1 |
| | Furcula furcula | GCA 911728495.1 |
| | Griposia aprilina | GCA 916610205.1 |
| | Habrosyne pyritoides | GCA 907165245.1 |
| | Hecatera dysodea | GCA 905332915.1 |
| | Hedya salicella | GCA 905404275.1 |
| | Hesperia comma | GCA 905404135.1 |
| | Hydriomena furcata | GCA 912999785.1 |
| | Hylaea fasciaria | GCA 905147375.1 |

| | | |
|---|---|---|
| | Idaea aversata | GCA 907269075.1 |
| | Laspeyria flexula | GCA 905147015.1 |
| | Leptidea sinapis | GCA 905404315.1 |
| | Limenitis camilla | GCA 905147385.1 |
| | Lysandra bellargus | GCA 905333045.1 |
| | Lysandra coridon | GCA 905220515.1 |
| | Mamestra brassicae | GCA 905163435.1 |
| | Marasmarcha lunaedactyla | GCA 923062675.1 |
| | Mesoligia furuncula | GCA 916614155.1 |
| | Mythimna impura | GCA 905147345.1 |
| | Noctua janthe | GCA 910589295.1 |
| | Ochropleura plecta | GCA 905475445.1 |
| | Ochlodes sylvanus | GCA 905404295.1 |
| | Omphaloscelis lunosa | GCA 916610215.1 |
| | Orgyia antiqua | GCA 916999025.1 |
| | Pammene fasciana | GCA 911728535.1 |
| | Papilio machaon | GCA 912999745.1 |
| | Peribatodes rhomboidaria | GCA 911728515.1 |
| | Pheosia gnoma | GCA 905404115.1 |
| | Philereme vetulata | GCA 918857605.1 |
| | Plebejus argus | GCA 905404155.1 |
| | Ptilodon capucinus | GCA 914767695.1 |
| | Schrankia costaestrigalis | GCA 905475405.1 |
| | Selenia dentaria | GCA 917880725.1 |
| | Sesia apiformis | GCA 914767545.1 |
| | Spilarctia lutea | GCA 916048165.1 |
| | Synanthedon vespiformis | GCA 918317495.1 |
| | Tinea semifulvella | GCA 910589645.1 |
| | Xestia c-nigrum | GCA 916618015.1 |
| | Ypsolopha scabrella | GCA 910592155.1 |
| | Zeuzera pyrina | GCA 907165235.1 |
| | Zygaena filipendulae | GCA 907165275.1 |
| | Nemoura dubitans | GCA 921293005.1 |
| | Ancistrocerus nigricornis | GCA 916049575.1 |
| | Andrena haemorrhoa | GCA 910592295.1 |
| | Athalia rosae | GCA 917208135.1 |
| | Bombus campestris | GCA 905333015.2 |
| | Bombus hypnorum | GCA 911387925.1 |
| | Bombus pascuorum | GCA 905332965.1 |
| | Bombus sylvestris | GCA 911622165.1 |
| | Bombus terrestris | GCA 910591885.1 |

| | | |
|---|---|---|
| | Dolichovespula media | GCA 911387685.1 |
| | Dolichovespula saxonica | GCA 911387935.1 |
| | Ectemnius continuus | GCA 910591665.1 |
| | Ectemnius lituratus | GCA 910593735.1 |
| | Lasioglossum morio | GCA 916610235.2 |
| | Mimumesa dahlbomi | GCA 917499265.1 |
| | Nomada fabriciana | GCA 907165295.1 |
| | Nysson spinosus | GCA 910591585.1 |
| | Haliclystus octoradiatus | GCA 916610825.1 |
| | Phorcus lineatus | GCA 921293015.1 |
| | Lasiommata megera | GCA 928268935.1 |
| | Agriphila tristella | GCA 928269145.1 |
| | Volucella inflata | GCA 928272305.1 |
| | Agonopterix arenella | GCA 927399405.1 |
| | Calamotropha paludella | GCA 927399485.1 |
| | Acleris emargana | GCA 927399475.1 |
| | Sarcophaga caerulescens | GCA 927399465.1 |
| | Macaria notata | GCA 927399415.1 |
| | Mythimna albipuncta | GCA 929112965.1 |
| | Agrypnus murinus | GCA 929113105.1 |
| | Limnephilus rhombicus | GCA 929108145.1 |
| | Andrena dorsata | GCA 929108735.1 |
| | Andrena minutula | GCA 929113495.1 |
| | Epistrophe grossulariae | GCA 929447395.1 |
| | Watsonalla binaria | GCA 929442735.1 |
| | Sarcophaga rosellei | GCA 930367235.1 |
| | Pollenia angustigena | GCA 930367215.1 |
| | Bombus pratorum | GCA 930367275.1 |
| | Myathropa florea | GCA 930367185.1 |
| | Opisthograptis luteolata | GCA 931315375.1 |
| | Plutella xylostella | GCA 932276165.1 |
| | Apotomis betuletana | GCA 932273695.1 |
| | Protocalliphora azurea | GCA 932274085.1 |
| | Leucozona laternaria | GCA 932273885.1 |
| | Diarsia rubi | GCA 932274075.1 |
| | Sarcophaga variegata | GCA 932273835.1 |
| | Patella vulgata | GCA 932274485.1 |
| | Podabrus alpinus | GCA 932274525.1 |
| | Epinotia nisella | GCA 932294315.1 |
| | Alitta virens | GCA 932294295.1 |
| | Chloroclysta siterata | GCA 932294275.1 |

| | | |
|---|---|---|
| | Agriopis marginaria | GCA 932305915.1 |
| | Pandemis cinnamomeana | GCA 932294345.1 |
| | Aporophyla lueneburgensis | GCA 932294355.1 |
| | Allophyes oxyacanthae | GCA 932294325.1 |
| | Diachrysia chrysitis | GCA 932294365.1 |
| | Meta bourneti | GCA 933210815.1 |
| | Bombylius major | GCA 932526495.1 |
| | Lobophora halterata | GCA 932526365.1 |
| | Ecliptopera silaceata | GCA 932527185.1 |
| | Philonthus cognatus | GCA 932526585.1 |
| | Caradrina clavipalpis | GCA 932526535.1 |
| | Phragmatobia fuliginosa | GCA 932526445.1 |
| | Nephrotoma flavescens | GCA 932526605.1 |
| | Epicampocera succincta | GCA 932526305.1 |
| | Nematostella vectensis | GCA 932526225.1 |
| | Operophtera brumata | GCA 932527175.1 |
| | Rhingia campestris | GCA 932526625.1 |
| | Lasioglossum pauxillum | GCA 933228785.1 |
| | Platycnemis pennipes | GCA 933228895.1 |
| | Machimus atricapillus | GCA 933228815.1 |
| | Miltochrista miniata | GCA 933228765.1 |
| | Hipparchia semele | GCA 933228805.1 |
| | Amblyteles armatorius | GCA 933228735.1 |
| | Leistus spinibarbis | GCA 933228885.1 |
| | Pemphredon lugubris | GCA 933228935.1 |
| | Stomorhina lunata | GCA 933228675.1 |
| | Buathra laborator | GCA 934046635.1 |
| | Leuctra nigra | GCA 934045905.1 |
| | Ypsolopha sequella | GCA 934047225.1 |
| | Apeira syringaria | GCA 934044485.1 |
| | Yponomeuta sedellus | GCA 934045075.1 |
| | Xylocampa areola | GCA 935421205.1 |
| | Polydrusus cervinus | GCA 935413205.1 |
| | Melolontha melolontha | GCA 935421215.1 |
| | Gibbula magus | GCA 936450465.1 |
| | Protodeltote pygarga | GCA 936450705.1 |
| | Rutpela maculata | GCA 936432065.1 |
| | Barbus barbus | GCA 936440315.1 |
| | Cheilosia pagana | GCA 936431705.1 |
| | Meganola albula | GCA 936450015.1 |
| | Glyphotaelius pellucidus | GCA 936435175.1 |

| | | |
|---|---|---|
| | Lepidonotus clava | GCA 936440205.1 |
| | Nowickia ferox | GCA 936439885.1 |
| | Synanthedon andrenaeformis | GCA 936446665.1 |
| | Sarcophaga subvicina | GCA 936449025.1 |
| | Hypsopygia costalis | GCA 937001555.1 |
| | Orcinus orca | GCA 937001465.1 |
| | Brachylomia viminalis | GCA 937001585.1 |
| | Bombylius discolor | GCA 939192795.1 |
| | Polyommatus icarus | GCA 937595015.1 |
| | Halyzia sedecimguttata | GCA 937662695.1 |
| | Aricia artaxerxes | GCA 937612035.1 |
| | Thecophora atra | GCA 937620795.1 |
| | Cistogaster globosa | GCA 937654795.1 |
| | Thera britannica | GCA 939531255.1 |
| | Metschnikowia zobellii | GCA 939531405.1 |
| | Trichoderma pseudokoningii | GCA 943193705.1 |
| | Xestia sexstrigata | GCA 941918905.1 |
| | Mucor piriformis | GCA 943193625.1 |
| | Phyto melanocephala | GCA 941918925.1 |
| | Sthenelais limicola | GCA 942159475.1 |
| | Calliphora vomitoria | GCA 942486065.1 |
| | Agrotis puta | GCA 943136025.1 |
| | Ophonus ardosiacus | GCA 943142095.1 |
| | Acrobasis suavella | GCA 943193695.1 |
| | Acentria ephemerella | GCA 943193645.1 |
| | Adalia bipunctata | GCA 910592335.1 |
| | Eupithecia abbreviata | GCA 943735965.1 |
| | Stelis phaeoptera | GCA 943735885.1 |
| | Sesia bembeciformis | GCA 943735995.1 |
| | Myopa tessellatipennis | GCA 943737955.1 |
| | Pherbina coryleti | GCA 943735915.1 |
| | Pollenia amentaria | GCA 943735925.1 |
| | Piscicola geometra | GCA 943735955.1 |
| | Tenthredo mesomela | GCA 943736025.1 |
| | Agriphila geniculea | GCA 943789525.1 |
| | Agaricus bisporus | GCA 943193715.1 |
| | Sphaerophoria taeniata | GCA 943590905.1 |
| | Nebria salina | GCA 944039245.1 |
| | Tiphia femorata | GCA 944319695.1 |
| | Rhogogaster chlorosoma | GCA 944452935.1 |
| | Stenoptilia bipunctidactyla | GCA 944452665.1 |

| | | |
|---|---|---|
| | Chrysolina oricalcia | GCA 944452925.1 |
| | Tachina lurida | GCA 944452675.1 |
| | Ophion luteus | GCA 944452715.1 |
| | Carterocephalus palaemon | GCA 944567765.1 |
| | Phosphuga atrata | GCA 944588485.1 |
| | Eupithecia centaureata | GCA 944547425.1 |
| | Micropterix aruncella | GCA 944548615.1 |
| | Euclidia mi | GCA 944739405.1 |
| | Andrena hattorfiana | GCA 944738655.1 |
| | Nebria brevicollis | GCA 944738965.1 |
| | Synanthedon myopaeformis | GCA 944738685.1 |
| | Eristalinus sepulchralis | GCA 944738645.1 |
| | Cryptosula pallasiana | GCA 945261195.1 |
| | Megachile willughbiella | GCA 945859595.1 |
| | Megachile ligniseca | GCA 945859555.1 |
| | Synanthedon formicaeformis | GCA 945859745.1 |
| | Anorthoa munda | GCA 945859665.1 |
| | Apamea sordens | GCA 945859715.1 |
| | Drepana falcataria | GCA 945859725.1 |
| | Lumbricus rubellus | GCA 945859605.1 |
| | Amphipoea oculea | GCA 945859645.1 |
| | Eupeodes corollae | GCA 945859685.1 |
| | Episyrphus balteatus | GCA 945859705.1 |
| | Epinotia demarniana | GCA 945867215.1 |
| | Flammulina velutipes | GCA 945909995.1 |
| | Isoperla grammatica | GCA 945910005.1 |
| | Brachypalpus laphriformis | GCA 945910035.1 |
| | Pleurotus ostreatus | GCA_947034855.1 |
| | Meconema thalassinum | GCA_946902985.1 |
| | Tridacna crocea | GCA 943736015.1 |
| | Tridacna gigas | GCA 945859785.1 |
| | Telmatherina bonti | GCA 933228915.1 |
| | Thunnus maccoyii | GCA 910596095.1 |
| | Thunnus albacares | GCA 914725855.1 |
| | Melinaea marsaeus rileyi | GCA 918358865.1 |
| | Limnoperna fortunei | GCA 944474755.1 |
| | Hyla sarda | s3://genomeark/species/Hyla_sarda/aHylSar1/assembly_MT_rockefeller/aHylSar1.MT.20220615.fasta |
| | Acridotheres tristis | s3://genomeark/species/Acridotheres_tristis/bAcrTri1/assembly_MT_rockefeller/bAcrTri1.MT.20220617.fasta |
| | Scomber japonicus | s3://genomeark/species/Scomber_japonicus/fScoJap1/assembly_MT_rockefeller/fScoJap1.MT.20220617.fasta |
| | Cynocephalus volans | s3://genomeark/species/Cynocephalus_volans/mCynVol1/assembly_MT_rockefeller/mCynVol1.MT.20220615.fasta |
| | Macropus eugenii | s3://genomeark/species/Macropus_eugenii/mMacEug1/assembly_MT_rockefeller/mMacEug1.MT.20220617.fasta |

| | | |
|---|---|---|
| | Mesoplodon densirostris | s3://genomeark/species/Mesoplodon_densirostris/mMesDen1/assembly_MT_rockefeller/mMesDen1.MT.20220617.fasta |
| | Monodelphis domestica | s3://genomeark/species/Monodelphis_domestica/mMonDom1/assembly_MT_rockefeller/mMonDom1.MT.20220617.fasta |
| | Neofelis nebulosa | s3://genomeark/species/Neofelis_nebulosa/mNeoNeb1/assembly_MT_rockefeller/mNeoNeb1.MT.20220615.fasta |
| | Nycticebus coucang | s3://genomeark/species/Nycticebus_coucang/mNycCou1/assembly_MT_rockefeller/mNycCou1.MT.20220615.fasta |
| | Sorex araneus | s3://genomeark/species/Sorex_araneus/mSorAra1/assembly_MT_rockefeller/mSorAra1.MT.20220617.fasta |
| | Hypanus sabinus | s3://genomeark/species/Hypanus_sabinus/sHypSab1/assembly_MT_rockefeller/mSorAra1.MT.20220617.fasta |
