## SupplementaryTable2 for "MitoHiFi: a python pipeline for mitochondrial genome assembly from PacBio High Fidelity reads"

| **Supplementary Table 2:** Species with previous mitochondrial genome assemblies found on INSC databates. Species in Figure 3A |
|---|
| Flammulina velutipes |
| Acipenser ruthenus |
| Agaricus bisporus |
| Plebejus argus |
| Canis lupus |
| Pleurotus ostreatus |
| Phragmatobia fuliginosa |
| Spilosoma lubricipeda |
| Neofelis nebulosa |
| Nycticebus coucang |
| Pararge aegeria |
| Pieris rapae |
| Thunnus maccoyii |
| Diadumene lineata |
| Barbus barbus |
| Scomber japonicus |
| Orcinus orca |
| Cervus elaphus |
| Mesoplodon densirostris |
| Aporia crataegi |
| Eristalis arbustorum |
| Syritta pipiens |
| Pyrgus malvae |
| Xestia c-nigrum |
| Thunnus albacares |
| Rattus norvegicus |

| |
|---|
| Melitaea cinxia |
| Operophtera brumata |
| Pholis gunnellus |
| Papilio machaon |
| Pieris napi |
| Euproctis similis |
| Melanostoma mellinum |
| Monodelphis domestica |
| Calliphora vomitoria |
| Mamestra brassicae |
| Eristalinus sepulchralis |
| Limenitis camilla |
| Ocypus olens |
| Episyrphus balteatus |
| Lumbricus rubellus |
| Piscicola geometra |
| Coccinella septempunctata |
| Sorex araneus |
| Accipiter gentilis |
| Limnoperna fortunei |
| Eristalis tenax |
| Lycaena phlaeas |
| Platycheirus albimanus |
| Plutella xylostella |
| Acridotheres tristis |
| Pterostichus madidus |
| Ischnura elegans |

| |
|---|
| Eupeodes latifasciatus |
| Nebria brevicollis |
| Eupeodes corollae |
| Tridacna crocea |
| Sarcophaga caerulescens |
| Tridacna gigas |
| Bombus terrestris |
