## Supplementary figures and images for "MitoHiFi: a python pipeline for mitochondrial genome assembly from PacBio High Fidelity reads"

### SupplementaryFigure2

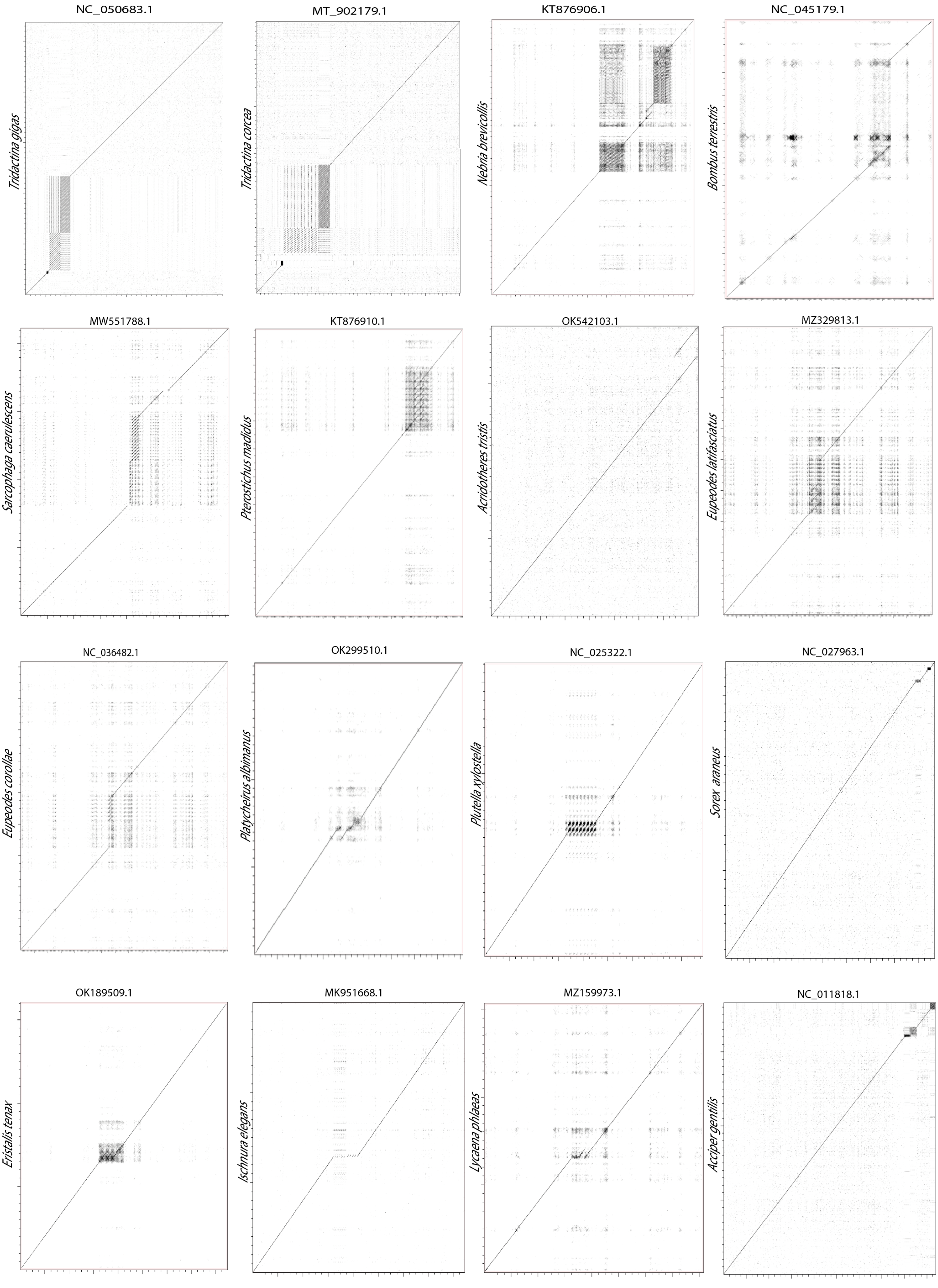
