## SupplementaryFigure1 for "MitoHiFi: a python pipeline for mitochondrial genome assembly from PacBio High Fidelity reads"

A

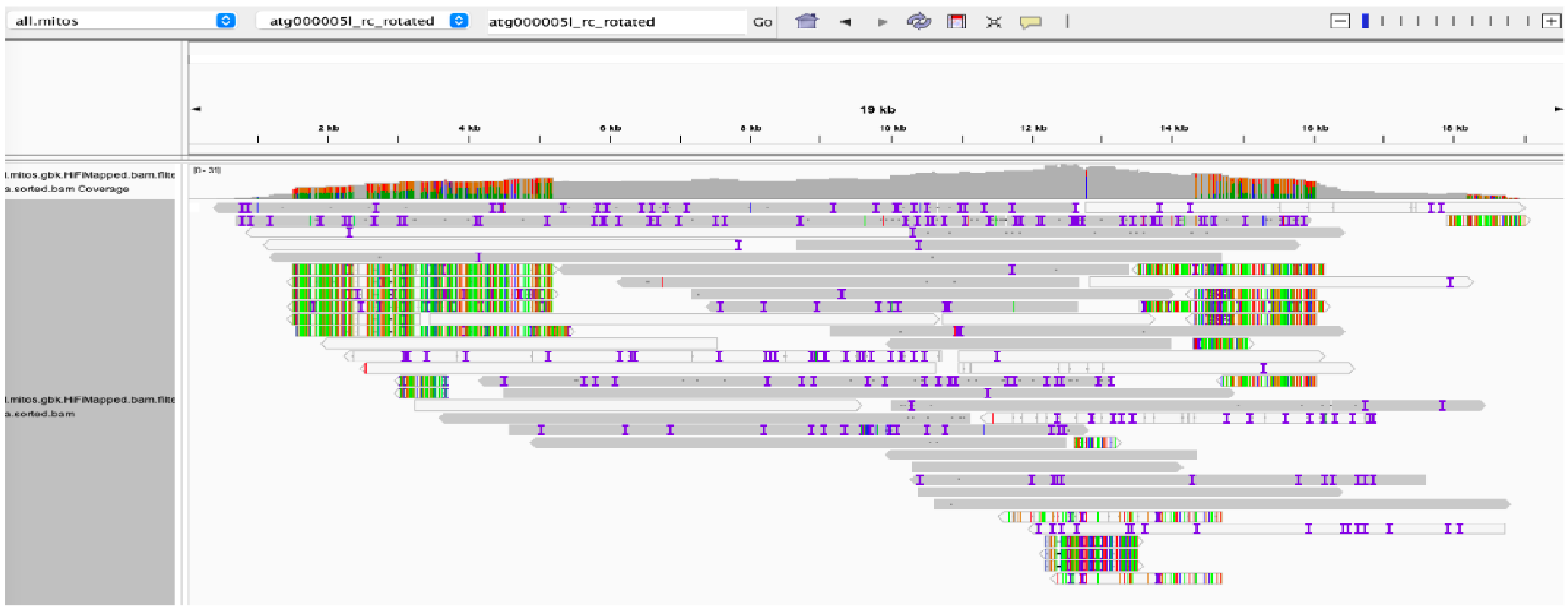


B

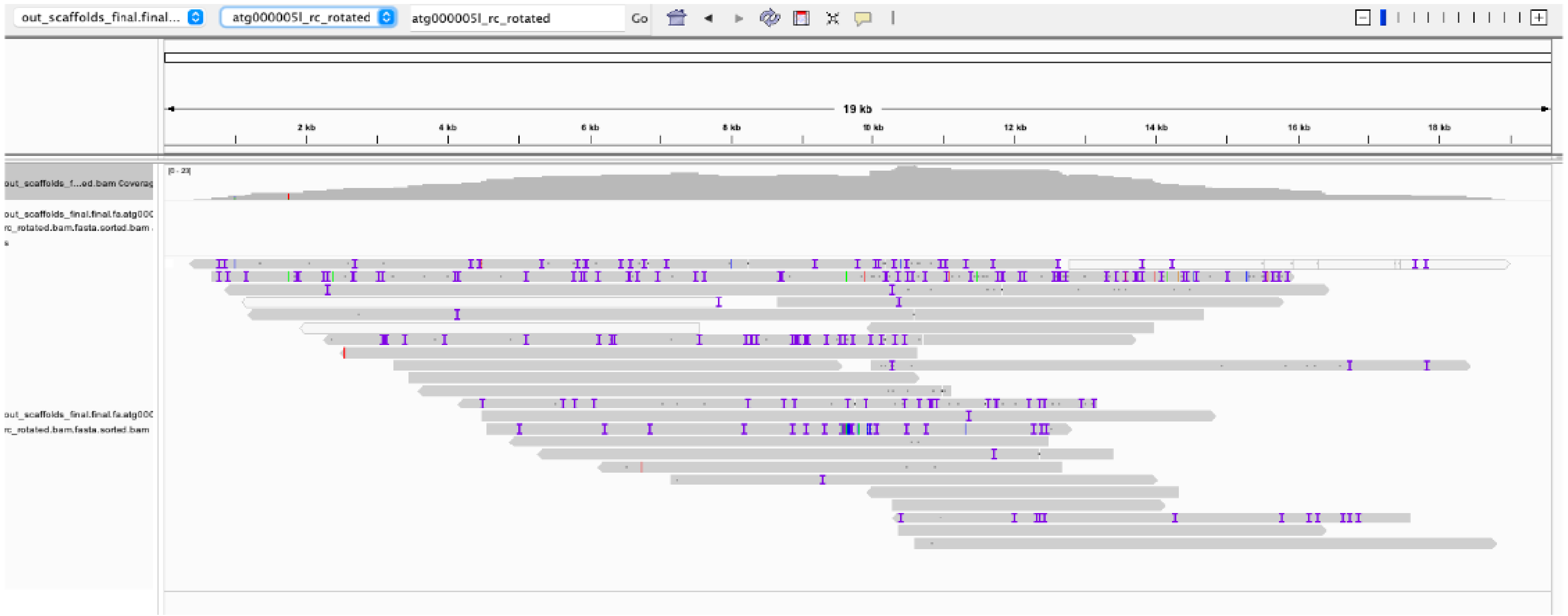


Supplementary Figure 2: Pannel A shows an IGV plot of all MitoHiFi filtered reads mapped back to the final_mitogenome.fasta of Andrena bucephala. In Pannel B, when reads are mapped to the final_mitogenome.fasta together with the nuclear genome, nUMTs reads are attracted to map at their nuclear locations and the variation track is gone.
